# Reduced Functional Coordination within the Default Mode Network in Schizophrenia During Naturalistic Neuroimaging

**DOI:** 10.64898/2026.08.11.744321

**Authors:** Yizhou Lyu, Yixuan Lisa Shen, Lourdes Concepcion Esparza, Eric A. Reavis, Carolyn Parkinson

## Abstract

**Background:** Social dysfunction is a major source of disability in schizophrenia, yet the neural mechanisms that contribute to impaired social understanding remain poorly understood. Converging evidence points to the role of the default mode network (DMN) in integrating social information over time to construct interpretations of social behaviors. Here, we tested the hypothesis that individuals with schizophrenia show reduced stimulus-driven coordination between brain regions within the DMN during free viewing of naturalistic social stimuli.

**Methods:** A sample of 124 adults (schizophrenia: *n*=63; healthy controls: *n*=61) viewed naturalistic video clips during fMRI. Inter-subject functional connectivity (ISFC) was computed within the two groups. Group differences were identified via permutation testing. We also explored group differences in other brain networks to examine whether effects were specific to the DMN.

**Results:** Individuals with schizophrenia showed weaker stimulus-driven coupling within the DMN compared to healthy controls, specifically between areas such as the parahippocampal gyrus, precuneus, and medial prefrontal cortex. Group differences in ISFC were specific to the DMN. Furthermore, no between-group differences emerged for within-participant functional connectivity in the DMN, suggesting that the observed effects reflect reduced stimulus-driven coordination among DMN regions when processing social stimuli rather than a more general decline in DMN connectivity.

**Conclusions:** Schizophrenia is characterized by impaired coordination within the DMN as it dynamically integrates social information over time, which could contribute to difficulties in constructing coherent interpretations of real-world social situations. These findings suggest that disrupted stimulus-driven network coordination might underlie social cognitive impairments in schizophrenia, highlighting the value of naturalistic paradigms for revealing network-level dysfunction under conditions that closely approximate real-world experience.

## Introduction

Individuals with a schizophrenia diagnosis often struggle with social information processing, including difficulties in perceiving, understanding, and navigating the social world (1,2). These challenges, broadly referred to as social cognitive deficits, include problems with accurately forming and maintaining emotional perceptions, inferring others’ mental states, and appraising social interactions (3–6). Social cognitive deficits in schizophrenia are linked to a variety of functional impairments, including difficulties in forming and maintaining social relationships, maintaining employment, and living independently (7,8). However, the neural mechanisms that give rise to these deficits remain poorly understood.

Schizophrenia has been associated with disrupted functioning of the default mode network (DMN), which is a distributed network integral to supporting social understanding (9,10). Resting-state fMRI studies have found reduced activation and connectivity among core DMN regions, such as the medial frontal and posterior cingulate cortices (2,11–13). Such reductions in DMN connectivity have been linked to social impairments, including worse social adjustment, perspective taking, and community functioning (14–16). Moreover, disrupted network activation patterns have been found to predict poorer social and functional outcomes (17,18). These findings suggest that reduced resting-state DMN connectivity may contribute to the social cognitive deficits observed in schizophrenia.

Despite extensive research showing that resting-state DMN connectivity is altered in schizophrenia and such alterations may contribute to social dysfunction, very little is known about DMN connectivity under real-world conditions in this population. Indeed, almost all studies of the DMN in psychosis have relied on resting-state scans or data from highly controlled tasks that bear little resemblance to real-life experiences of the social world. In the present study, we addressed this knowledge gap by adopting a more naturalistic video-watching task in which viewers dynamically integrate information from multiple sensory modalities, interpret characters’ motives, and draw on prior knowledge to understand hidden meanings in the social narratives (19,20).

To study how DMN regions work together to process information about complex social narratives in schizophrenia, we used a method that measures the stimulus-driven coordination among regions of the DMN: inter-subject functional connectivity (ISFC). Unlike traditional functional connectivity (FC) measures that assess the extent to which brain activity covaries over time across brain regions within individuals (21), ISFC assesses the extent to which activity in one brain region covaries with activity in another region *across* individuals. Specifically, ISFC is computed by correlating the response time series from a given brain region in one person with the response time series of another brain region, averaged across other participants, during periods when all participants are experiencing the same stimuli (22). As such, ISFC is closely related to inter-subject correlation (ISC), a complementary method that focuses on within-region synchrony of brain activity across individuals, rather than inter-regional coordination.

A major advantage of the ISFC method over traditional FC approaches is that ISFC filters out sources of noise which can be mistaken for signal in traditional FC analyses, such as head motion or physiological drift. More importantly for the purposes of the current study, ISFC isolates *stimulus-driven* neural fluctuations, filtering out unrelated fluctuations due to intrinsic brain activity that is not time-locked to the stimulus (23,24). Using ISFC, previous work in healthy samples has found that regions of the DMN dynamically strengthen their coupling as listeners construct and update the meaning of an unfolding story, and that greater stimulus-driven coupling between regions of the DMN (i.e., greater ISFC) corresponds to more effective encoding of the narrative (22). Taken together, these findings suggest that stimulus-driven DMN connectivity may provide a sensitive index of high-level social information processing and narrative comprehension, and thus may be helpful for understanding sociocognitive impairments in schizophrenia.

In the present study, we examined stimulus-driven coordination among regions of the DMN in participants with schizophrenia and healthy controls as they watched video clips in the fMRI scanner. We tested whether the DMN showed differences in stimulus-driven inter-regional coupling during naturalistic narrative processing across groups. We predicted that participants with schizophrenia would exhibit reduced ISFC within the DMN, reflecting weaker coordination among DMN regions during the processing of complex social narratives. This may reflect weaker engagement of a core network that supports the integration and interpretation of socially relevant information, providing a potential neural mechanism underlying the social cognitive impairments observed in schizophrenia.

## Methods and Materials

### Participants

The present dataset included an interim sample of 124 individuals from an ongoing cross-sectional study, including 61 healthy controls and 63 participants diagnosed with schizophrenia. For recruitment details, see Y.L. Shen et al. (25).

Each participant completed a series of clinical interviews and self-report measures to confirm eligibility. Structured diagnostic interviews (SCID-I) were administered to establish or rule out schizophrenia, and healthy controls additionally completed the SCID-PD (26). Clinical ratings, including the Expanded Brief Psychiatric Rating Scale (BPRS) (27), the Clinical Assessment Interview for Negative Symptoms (CAINS) (5), and the Hamilton Depression Scale (28), were used to rate clinical symptoms. Participants also provided demographic information and information about current medications and medical history.

All clinical interviewers received training from the Treatment Unit of the Greater Los Angeles VA VISN 22 Mental Illness Research Education and Clinical Center and followed a standardized protocol. All study activities were approved by the UCLA Institutional Review Board (IRB-21-1219), and written informed consent was obtained from each individual. All participants were reimbursed for their time.

### Experimental procedure

#### fMRI data acquisition

All fMRI scans were performed on a 3T Siemens Prisma system with a 32-channel head coil at the UCLA Staglin Center for Cognitive Neuroscience. The session began with a short localizer scan, followed by a high-resolution T1-weighted magnetization-prepared rapid gradient-echo (MPRAGE) acquisition (0.8 mm isotropic voxels). Field maps were acquired to compensate for geometric distortions caused by magnetic field inhomogeneities. Task-based functional data were obtained using a T2*-weighted gradient-echo echo planar imaging (EPI) sequence with a repetition time (TR) of 1000 ms, echo time (TE) of 36 ms, flip angle of 70°, multiband factor of 6, a 104 × 104 voxel matrix, a field of view (FoV) of 208 mm, and an isotropic voxel size of 2 mm. During each run, participants viewed visual stimuli on an MR-compatible LCD screen (Cambridge Research Systems, Rochester, UK) via MATLAB, and audio was delivered through noise-canceling headphones (Optoacoustic Ltd., Mazor, Israel).

#### fMRI Tasks

All participants underwent a single fMRI session. During the scan, they viewed eight video clips presented in a fixed order across three runs (Run 1: 15 min 32 s; Run 2: 13 min 40 s; Run 3: 11 min 26 s). The clips varied widely in style and content, with the intention of eliciting diverse cognitive and affective responses. During each run, participants were instructed to watch these clips as they would if casually viewing television or online videos, and to remain as still as possible.

### Data analysis

#### fMRI preprocessing and parcellation

We preprocessed the fMRI data using *fMRIPrep* (version 22.0.2) (29) using default parameters. Briefly, T1-weighted images were bias-field corrected, skull-stripped, tissue-segmented, and nonlinearly normalized to the MNI152Nlin2009cAsym template. For each BOLD run, a skull-stripped EPI reference volume was co-registered to the T1-weighted image using boundary-based registration; head motion was estimated and corrected; and standard confound time series were generated. After fMRIPrep preprocessing, we spatially smoothed the data (6 mm FWHM kernel), regressed out nuisance signals (framewise displacement, global brain signals, and 6 motion parameters, CSF, white matter), and applied a 0.01Hz-0.08 Hz band-pass filter to mitigate low-frequency drift and high-frequency noise.

Parcellation of the brain was performed using the Shen 268-node atlas (30), which divides the brain into 268 non-overlapping regions, and can be organized into eight major networks: medial frontal, frontoparietal, default mode, subcortical-cerebellar, motor, visual I, visual II, and visual association. Our primary analyses used only the time series from the 20 parcels in the DMN as regions of interest (ROIs; **see Fig. 1**). For each participant’s three movie-viewing runs, the first fourteen time points of each run (corresponding to an 8-second fixation screen that precedes the videos and the subsequent 6 seconds to account for the hemodynamic lag of the BOLD response) were discarded to minimize activity unrelated to the videos.

**Figure 1.**
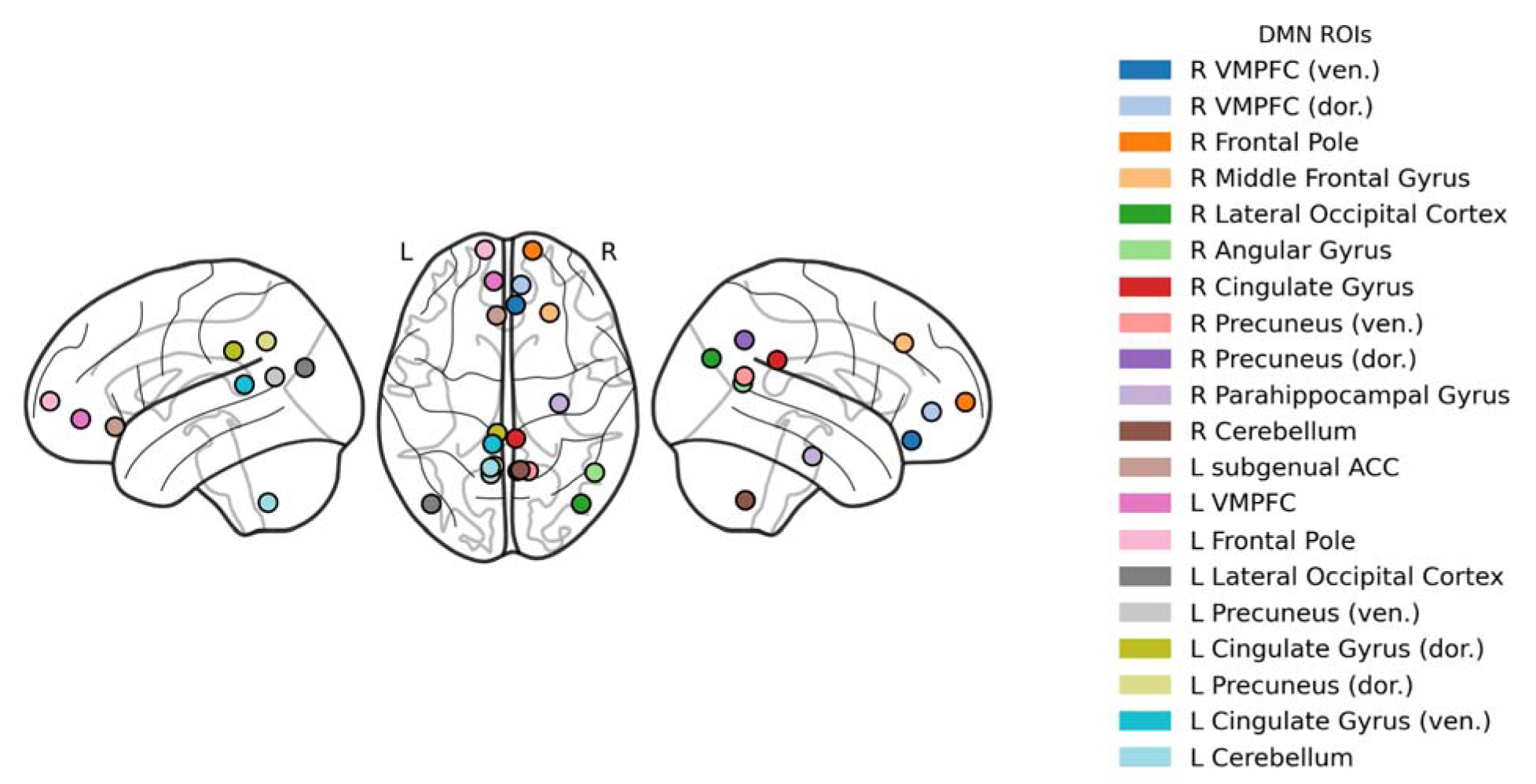
Shen 268-regions atlas. Center locations for all regions of interest in the DMN, defined by the Shen 268-region atlas, labelled and displayed in lateral (left/right) and dorsal views.

Motion quality control was applied run-by-run: at every TR, we computed the Euclidean norm of the three translational motion parameters and flagged volumes exceeding 3 mm. If more than 25% of TRs in a given run were flagged to have excessive motion, the entire run was excluded for the participant. Using this criterion, 13 participants in the schizophrenia group and 3 participants in the healthy control group had one or more runs excluded. After excluding high-motion runs, group differences in framewise displacement were not significant in runs 1 (*M_HC_*= 0.286 ± 0.167; *M_SZ_* = 0.336 ± 0.135; Welch’s *t*(108.74) = -1.73, *p* = 0.087) or 2 (*M_HC_* = 0.302 ± 0.191; *M_SZ_* = 0.342 ± 0.143; Welch’s *t*(105.52) = -1.27, *p* = 0.208). Run 3 showed a small but significant difference (*M_HC_* = 0.293 ± 0.168; *M_SZ_* = 0.373 ± 0.170; Welch’s *t*(113.95) = -2.55, *p* = 0.012).

#### Comparing Stimulus-Driven Coupling Among DMN Regions between Groups

For each participant, only time-series data from the 20 DMN ROIs were extracted, producing a 2D array (T time points × 20 DMN nodes). The time series for every ROI were *z*-scored within each participant to remove mean differences and scale variability, then concatenated across runs. Finally, a mean time series for each ROI was calculated by averaging across all voxels within the parcel, yielding a single time series for each ROI and participant.

We computed ISFC separately for the schizophrenia and healthy control groups. For each participant, we correlated the mean time series for each DMN ROI with the average time series of all other participants in the same group for every other DMN ROI. We symmetrized every participant’s ISFC matrix by averaging the corresponding upper-and lower-triangular cells. We also discarded the diagonal, as those cells reflect inter-subject correlations (i.e., effects *within* an ROI that do not reflect functional connectivity). This leaves an ISFC matrix comprised of 190 off-diagonal cells. When a participant had one or more runs excluded for motion, that participant’s ISFC calculation used group-average data constructed only from the included runs for that participant, ensuring the same temporal coverage for the individual and the group average. We then calculated a mean ISFC matrix for each group by averaging across all participants’ ISFC matrices within groups (see **Fig. 2**). Each off-diagonal cell in each participant’s ISFC matrix corresponds to the similarity of that participant’s response time series in one region of the DMN with that of other participants within their group in a different region of the DMN–i.e., region-to-region stimulus-driven functional connectivity.

**Figure 2.**
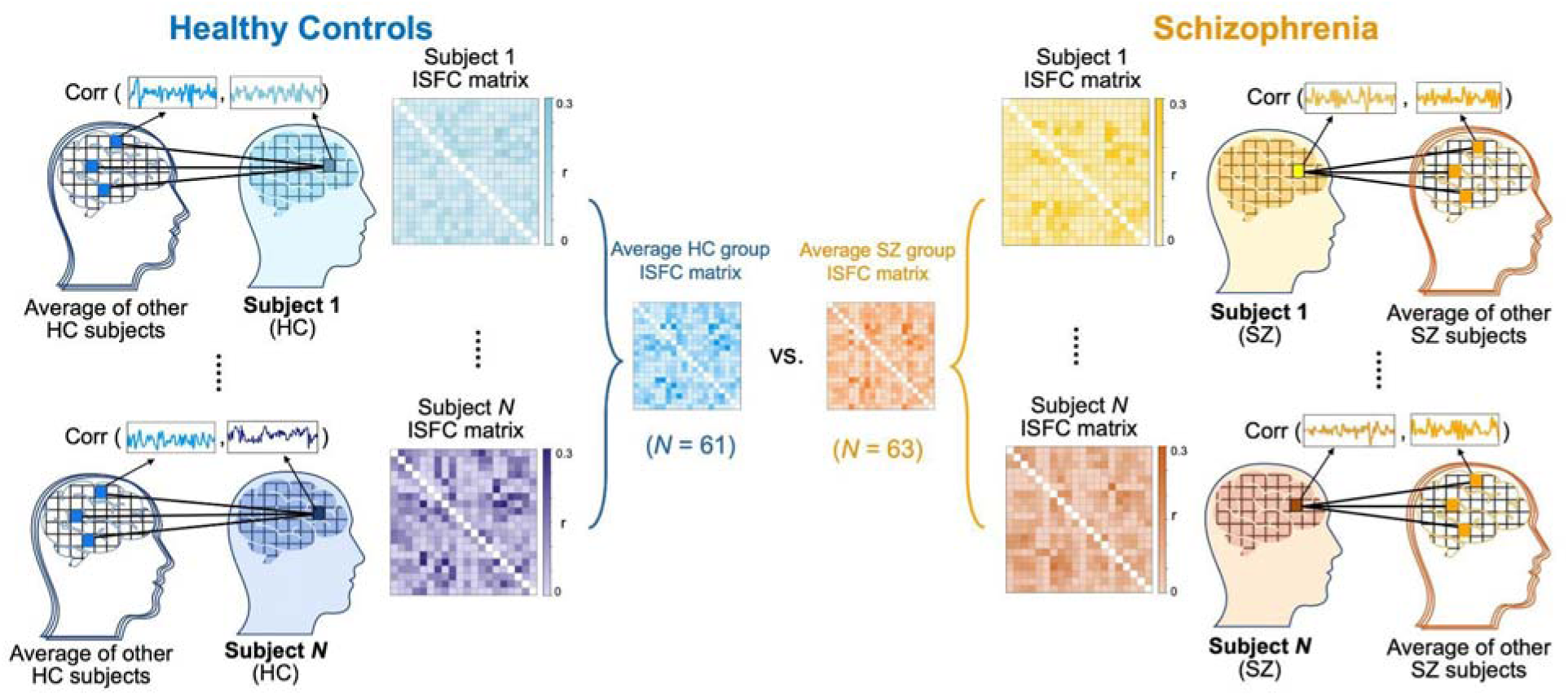
Schematic illustrating the ISFC method. For each region of interest (ROI) in the DMN, we correlated each subject’s ROI time series with the leave-one-out average across other members of the group for every other ROI to obtain a 20×20 matrix. Individual matrices were then averaged across subjects to yield a group ISFC. By correlating each participant’s activity with that of other participants while viewing the same stimulus, ISFC captures shared, stimulus-driven coupling between brain regions and disregards the influence of intrinsic neural activity that is not time-locked to the stimulus.

We next used permutation testing to identify significant ISFC differences between the two groups. For each of the 190 cells in the ISFC matrix, we computed the absolute difference in ISFC values between the two groups. We then generated a null distribution by randomly reassigning healthy control or schizophrenia patients’ labels (preserving the original group sizes) and recalculating absolute difference scores on the randomly labeled data, with 10,000 iterations. The likelihood of the observed differences occurring by chance (i.e., *p*-values) was computed as the proportion of permuted differences that were at least as extreme as the observed difference. We corrected our threshold for inferring significant group differences for the number of ISFC cells compared using a false discovery rate (FDR) procedure at *q* = 0.05 (31).

Following these primary analyses, we repeated the same procedure within the other seven networks defined by the Shen 268-regions atlas (30) to examine whether group differences in stimulus-driven functional connectivity were specific to the DMN. To emulate the stringency of our primary analyses, we corrected for multiple comparisons according to the number of edges within each network.

We also conducted a follow-up analysis to determine whether any observed group differences in ISFC might be explained by group differences in within-subject FC. For each participant, within each of the 8 networks, we computed an FC matrix by correlating the *z*-scored time series of each ROI with the time series of every other ROI within the *same* participant. The diagonal was discarded, and because FC matrices are symmetric, we retained the unique off-diagonal cells of the FC matrix. We then used the same FDR-corrected permutation-testing approach from our primary analyses to identify significant group differences in FC.

## Results

The schizophrenia and control participants were comparable in age, sex, race, and parental education (all *p*s > .26). The sample skewed male, reflecting a high proportion of Veteran participants in both groups. As expected, individuals with schizophrenia had significantly fewer years of education than controls (**Table 1**; *t*(118) = 2.85, *p* = .005). The schizophrenia sample consisted of chronic, stable outpatients who were not highly symptomatic at the time of testing, as reflected in relatively low mean BPRS positive symptom scores (*M* = 2.91, *SD* = 0.91) and CAINS Motivation and Pleasure (*M* = 1.57, *SD* = 0.70) and Expressive (*M* = 0.72, *SD* = 0.74) scores.

**Table 1.** Participant demographics.

| Variables | Healthy controls<br>(N = 61) | Schizophrenia<br>(N = 63) | Group comparison |
| --- | --- | --- | --- |
| Age, mean (SD) | 45.80<br>(13.06) | 47.41<br>(11.01) | $t(122) = -0.74$<br>($p = .459$) |
| Sex, $n$ (%) | | | $\chi^2(1, N = 124) = 1.23$<br>($p = .267$) |
| Male | 48 (78.7%) | 43 (68.3%) |  |
| Female | 13 (21.3%) | 20 (31.7%) |  |
| Race, $n$ (%) | | | $\chi^2(4, N = 124) = 1.35$<br>($p = .853$) |
| White | 26 (42.6%) | 25 (39.7%) |  |
| Asian | 7 (11.5%) | 5 (7.9%) |  |
| Black/African American | 17 (27.9%) | 22 (34.9%) |  |
| American Indian/Alaska Native | 1 (1.6%) | 2 (3.2%) |  |
| Other (more than 1 race) | 10 (16.4%) | 9 (14.3%) |  |
| Years of education (SD) | 14.72 (2.04) | 13.60 (2.27) | $t(118) = 2.85$<br>($p = .005$) |
| Years of parental education (SD) | 14.21 (3.03) | 13.55 (3.24) | $t(114) = 1.13$<br>($p = .262$) |

### Weaker stimulus-driven coupling between regions of the DMN in schizophrenia patients during social narrative perception

Stimulus-driven coordination (i.e., ISFC) among regions of the DMN was significantly different between groups, with 10 of the 190 within-DMN connections showing significantly greater ISFC in the healthy control group than in the schizophrenia group after FDR correction (**Fig. 3**). Specifically, anterior-posterior connections between regions of the frontal, temporal, and parietal lobes were significantly stronger in healthy controls than schizophrenia patients, especially between the parahippocampal gyrus (PHG), precuneus, and the medial prefrontal cortex (see **Fig. 3 and Table S1**). These connections were between the right parahippocampal gyrus and the right and left ventral medial prefrontal cortex (VMPFC), the right and left ventral precuneus, and the left subgenual anterior cingulate cortex (sgACC); between the left sgACC and the right VMPFC, as well as the right and left ventral precuneus; between the left and right VMPFC; and between the left and right ventral precuneus.

**Figure 3.**
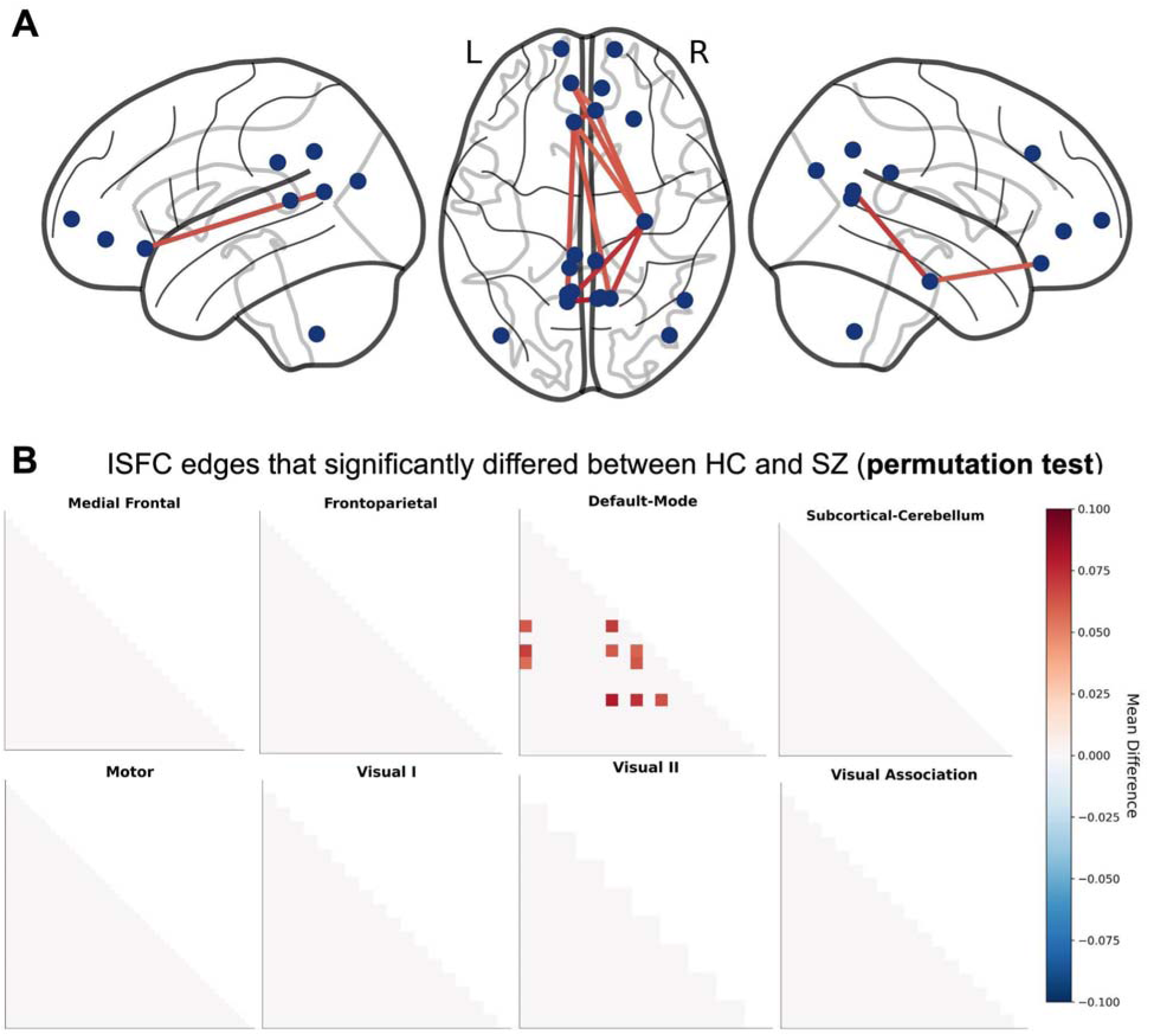
Group differences in stimulus-driven functional connectivity between areas of the default-mode network. **A.** Differences in mean ISFC values between schizophrenia and healthy control groups for the 190 within-DMN connections visualized on a glass brain, plotted only when the edge survived the group-label permutation test and FDR correction. **B.** Group difference in stimulus-driven coordination (ISFC) between regions within the eight networks of the Shen atlas. For each network, each within-network ISFC edge was compared between the healthy control and schizophrenia groups using 10,000 label-permutation tests that preserve the original group sizes; resulting *p*-values were FDR-corrected using the Benjamini-Hochberg method across all edges within that network.

### Weaker stimulus-driven coupling among regions is specific to the DMN

To explore possible stimulus-driven coupling within other brain networks, we conducted follow-up analyses in other brain networks. Specifically, we repeated the ISFC procedure for every other network defined in the Shen atlas. No significant differences in ISFC were found in any of the other seven networks (see **Fig. 3B**).

### Group differences in DMN ISFC do not recapitulate within-person FC effects

To test whether reduced ISFC in schizophrenia could simply reflect group differences in within-person FC, we compared FC between participants in the healthy control and schizophrenia groups within each of the eight Shen networks, including the DMN, using an analogous statistical procedure. Of the eight networks, only a few connections within the visual association network showed significant group differences in FC after correction (see **Fig. 4; Table S2**).

**Figure 4.**
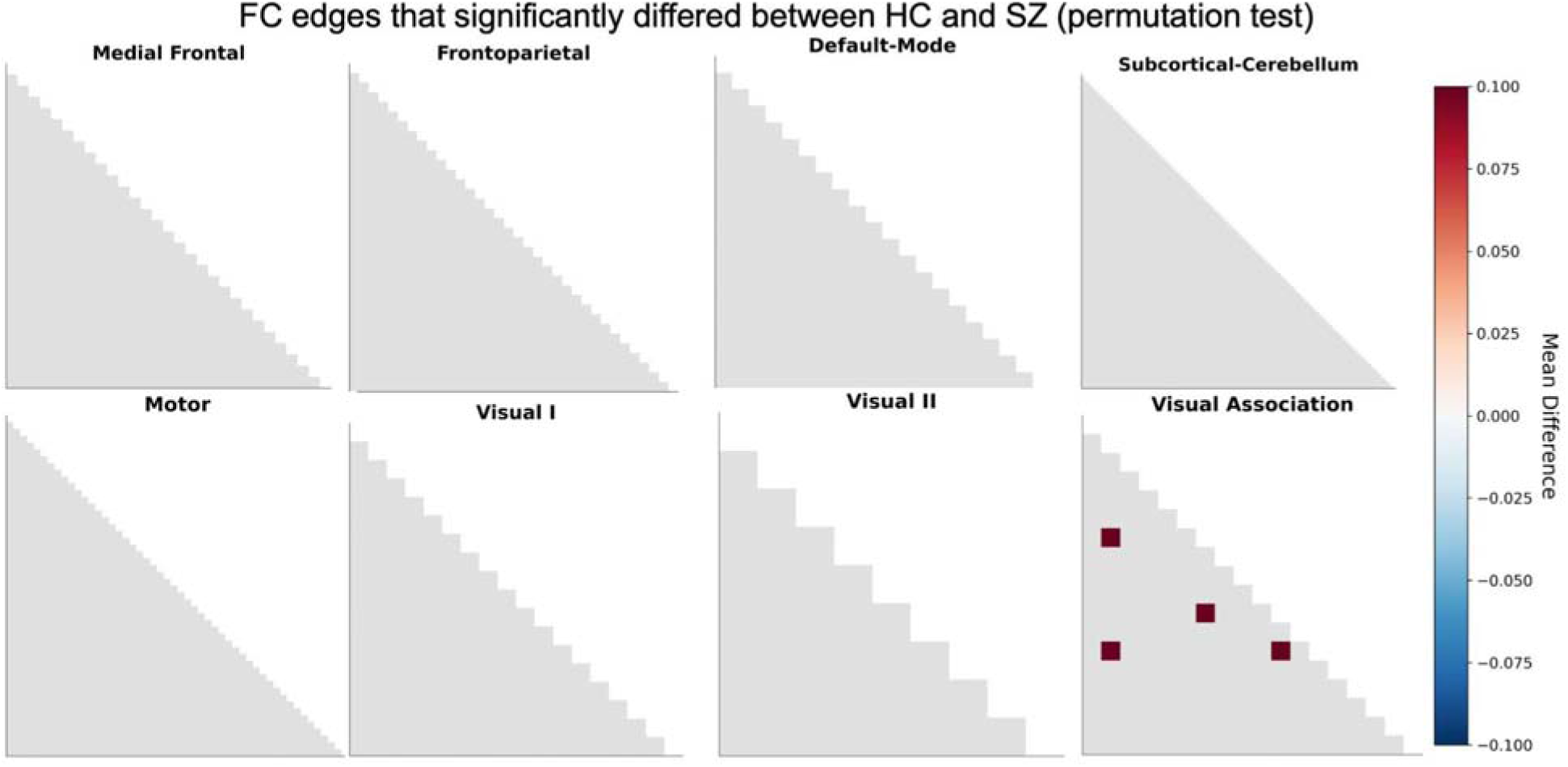
Group differences in functional connectivity across the eight Shen networks. For each network, each within-network FC edge was compared between the healthy control and schizophrenia groups using 10,000 label-permutation tests that preserve the original group sizes; resulting *p*-values were FDR-corrected using the Benjamini-Hochberg method across all edges within that network.

## Discussion

In this study, we found that individuals with schizophrenia exhibited reduced stimulus-driven coordination among regions of the DMN during naturalistic social processing. This disruption was specific to the DMN and was not observed in other large-scale brain networks. Moreover, we did not find group differences within the DMN using more traditional within-person FC analyses. This suggests that, in the context of naturalistic stimulation (here, viewing audiovisual videos that depict social narratives), schizophrenia may be characterized by impaired stimulus-driven coordination among DMN regions, which is distinct from the differences in intrinsic connectivity that are assessed in typical resting-state studies. Overall, the results suggest that schizophrenia is characterized by a disruption in how the DMN dynamically integrates information over time under conditions that approximate real-world social experience.

The DMN is thought to support high-level integration of information over time to support understanding of social narratives, including tracking others’ mental states, goals, and relationships across unfolding events (22,24). Prior work using ISC approaches (which examines inter-subject alignment *within* corresponding brain regions, in contrast to ISFC, which uses the inter-subject approach to instead capture stimulus-driven coupling *between* brain regions) has shown that activity within DMN regions becomes aligned across individuals during narrative processing (23,24), reflecting shared interpretation of the stimulus. Recent work in schizophrenia has reported reduced ISCs in these regions, suggesting less consistent stimulus-driven processing (25,32,33). However, the ISC approach captures synchrony within regions and does not examine how regions interact with one another. By using an ISFC approach, this study builds on prior findings, showing that schizophrenia is characterized by reduced stimulus-driven coupling between pairs of DMN regions during social narrative understanding. This implies a failure of the DMN to function as a coordinated network during naturalistic social processing.

While previous resting-state studies using within-person FC have found altered connectivity in the DMN and frontoparietal network in schizophrenia (e.g., 34,35), we did not find analogous group differences. This discrepancy is consistent with the expectation that naturalistic paradigms emphasize externally driven neural dynamics which may not be captured at rest. More specifically, prior studies that found FC effects in resting-state data presumably captured differences in intrinsically-driven coactivation patterns among regions, whereas movie-watching paradigms induce extrinsic, stimulus-driven coactivation patterns that could overshadow intrinsic coupling. Although some group differences in within-person FC were observed in the visual association network, such effects were limited and likely reflect lower-level perceptual processing differences in schizophrenia (36–39), rather than the higher-order social-cognitive processes thought to be indexed by DMN coupling.

A key direction for future research is to determine whether stimulus-driven connectivity patterns can serve as clinically meaningful biomarkers that predict variability in real-world outcomes or sensitivity to interventions. Because ISFC can capture dynamic, stimulus-driven coordination among brain regions under conditions that resemble real-world experiences, it could provide a sensitive index of neural processes that support social functioning in everyday life. Establishing links between ISFC and real-world outcomes could help inform diagnosis, prognosis, and intervention efforts in schizophrenia. Accordingly, we plan to examine the link between ISFC and real-world functional outcomes once data collection is finished.

While informative, the current findings are subject to several key limitations. First, while we applied multiple strategies to mitigate motion-related confounds, a small group difference in motion was observed in one run. However, given that a significant motion difference was found in only one of the three fMRI runs and that stringent mitigation strategies were used, it is unlikely that motion alone accounts for the present findings. Second, the current analyses focus on group-level differences and do not address the variability within the schizophrenia population. Individuals with schizophrenia vary widely in the severity and nature of the social cognitive impairments and related functional challenges they experience (7). Future work should examine whether reduced ISFC within the DMN when processing social narratives predicts individual differences in symptom profiles, social functioning, or treatment response among schizophrenia patients. Relatedly, the schizophrenia sample in this study consisted of chronic, clinically stable, medicated outpatients, which may limit generalizability to first-episode or more acutely symptomatic samples, as prior studies have found differences in FC across illness stages (40).

In summary, the present study demonstrates that reduced stimulus-driven coordination within the DMN occurs in schizophrenia during naturalistic social processing. These findings suggest that social cognitive impairments in schizophrenia may relate to a failure of distributed sets of brain regions to dynamically integrate information over time. More broadly, this work highlights the importance of examining brain function under ecologically valid conditions, showing that naturalistic stimuli, combined with analytic approaches that isolate stimulus-driven connectivity patterns, can reveal network-level dysfunction that might otherwise be obscured by traditional methods. By capturing how the brain coordinates activity in response to rich, dynamic stimuli that capture elements of real-world social experience, this approach offers a promising avenue for advancing understanding of the neural mechanisms that underlie social cognitive dysfunction in schizophrenia and other clinical populations.

## Acknowledgements

This project was funded by the National Institutes of Health (R01 MH128720; awarded to EAR and CP). Participant recruitment was supported by the Biomedical Informatics Program (BIP) within the UCLA Clinical and Translational Science Institute (NIH UL1TR001881). Computational and data-storage resources were provided by the Hoffman2 Cluster, operated by the Research Technology Group within the UCLA Office of Advanced Research Computing. The study sponsors were not involved in the conduct of the study.

Preliminary findings from this project were presented at the annual meeting of the Social and Affective Neuroscience Society. These data have not previously been published.

The authors acknowledge James Lopez, Melodie Yen, Yasmeen Campos, Jake Isenman, Gina Jackson, and Noah Moreno for their assistance with data collection.

