## Supplemental Table for "Reduced Functional Coordination within the Default Mode Network in Schizophrenia During Naturalistic Neuroimaging"

### **Supplementary Data**

**Table S1. Default Mode Network inter-subject functional connectivity edges significantly stronger in healthy control than schizophrenia patients**

|  | **Node 1** | **Node 2** | **Mean difference** |
| --- | --- | --- | --- |
| 1 | Right parahippocampal gyrus | Right ventral medial prefrontal cortex | 0.0611 |
| 2 | Right parahippocampal gyrus | Right ventral precuneus | 0.0698 |
| 3 | Left subgenual ACC | Right ventral medial prefrontal cortex | 0.0682 |
| 4 | Left subgenual ACC | Right ventral precuneus | 0.0610 |
| 5 | Left subgenual ACC | Right parahippocampal gyrus | 0.0589 |
| 6 | Left ventral medial prefrontal cortex | Right ventral medial prefrontal cortex | 0.0561 |
| 7 | Left ventral medial prefrontal cortex | Right parahippocampal gyrus | 0.0621 |
| 8 | Left ventral precuneus | Right ventral precuneus | 0.0791 |
| 9 | Left ventral precuneus | Right parahippocampal gyrus | 0.0727 |
| 10 | Left ventral precuneus | Left subgenual ACC | 0.0648 |

**Table S2. Visual association network functional connectivity edges significantly stronger in healthy control than schizophrenia patients**

|  | **Node 1** | **Node 2** | **Mean difference** |
| --- | --- | --- | --- |
| 1 | Temporal occipital fusiform cortex | Right lateral occipital cortex | 0.1324 |
| 2 | Inferior temporal gyrus | Right lateral occipital cortex | 0.1064 |
| 3 | Temporal occipital fusiform cortex | Left lateral occipital cortex | 0.1168 |
| 4 | Inferior temporal gyrus | Left lateral occipital cortex | 0.1060 |
